# Development and Utilization of 3D Anatomy Education Content using Metaverse and XR for Remote Telemedicine Education

**DOI:** 10.1101/2024.06.05.597650

**Authors:** Dong Hyeok Choi, Seo Yi Choi, So Hyun Ahn, Rena Lee, Sung Ho Cho, Seung Ho Han

**Author notes:** These authors contributed equally to this study (Dong Hyeok Choi, Seo Yi Choi). Correspondence to*: So Hyun Ahn, Seung Ho Han, and Rena Lee.

## Abstract

The objective of this study is to explore innovative integration within the field of anatomy education by leveraging HoloLens 2 Augmented Reality Head-Mounted Display (AR HMD) technology and real-time cloud rendering. Initial 3D datasets, comprising extensive anatomical information for each bone, were obtained through the 3D scanning of a full-body cadaver of Korean male origin. Subsequently, these datasets underwent refinement processes aimed at enhancing visual fidelity and optimizing polygon counts, utilizing Blender software. Unity was employed for the development of the Metaverse platform, incorporating tailored 3D User Experience (UX) and User Interface (UI) components to facilitate interactive anatomy education via imported cadaver models. Integration with real-time remote rendering cloud servers, such as Azure, was implemented to augment the performance and rendering capabilities of the HoloLens 2 AR HMD. The extended reality (XR) content uses the Photon Cloud network for real-time data synchronization and HoloLens 2 voice functionality. The metaverse platform supports user interaction through room creation and joining, with various tools for bone manipulation, color differentiation, and surface output. Collaboration features enable sharing and synchronization of model states. The study highlights the importance of technological innovation in anatomy education for future medical professionals. The proposed content aims to address limitations of traditional methods and enhance learning experiences. Continued efforts in developing and improving such technologies are crucial to equip learners with essential skills for adaptation in the evolving healthcare landscape. keyword: Metaverse, Anatomy, Cadaver, Extended Reality(XR), Virtual Reality(VR)

## INTRODUCTION

Anatomy education plays a crucial role in medical training by providing a foundational understanding of the human anatomy (1). Anatomy education plays a fundamental component not only for aspiring surgical doctors but also for students in nursing departments and other healthcare fields (2, 3). However, traditional methods of teaching anatomy, such as cadaver dissection, have raised ethical concerns, financial constraints, and other limitations. Cadavers are not always accessible as there are challenges in specimen availability, preservation, maintenance, as well as the high cost of implementation (4–6). The declining trend in cadaver use raises concerns about the reduced opportunities for aspiring medical students to gain practical experience in dissection and gross anatomy (7, 8). Social distancing and restrictions as responses to pandemic situations, like the recent COVID-19 pandemic, has further exacerbated this issue, leading to a significant reduction in the use of cadavers in anatomy education. A survey conducted among anatomy educators in the United States has shown a decline in using cadavers, specifically 39% in gross anatomy courses and 43% in other anatomy courses during the pandemic (9).

In time, advancements in technology have provided opportunities to develop integrative approaches that address ethical concerns and have helped to improve the level of comprehensiveness in anatomy education for students (10, 11). The use of imaging technologies such as MRI and CT scans has enabled researchers to visualize the internal structures of the human body with unprecedented detail. Additionally, virtual reality (VR) and augmented reality (AR) has been used to create interactive 3D models that allow students to explore the body and its systems in an engaging and immersive way. The integration of these technologies has revolutionized the delivery of medical education and expanded the possibilities for remote learning and digital resources (12–18).

The effects of COVID-19 have even accelerated the need for integrating the latest interactive technology in medical education, highlighting the importance of innovative solutions such as extended reality (XR) technology and the metaverse. The use of XR and the metaverse in clinical practice education is gaining popularity in addressing these challenges (19). For example, the University of Northampton is using VR technology to provide nursing education (20), while the University of Newcastle is using VR simulations for childbirth education (21). As such, the integration of XR technology in medical education offers numerous advantages, such as enhancing student learning outcomes, providing hands-on experience within a safe simulated environment, promoting interprofessional education, saving costs and time, and the potential to revolutionize the way medical education is delivered (22).

Although the integration of XR technology and attempts in anatomy education offer numerous advantages, several challenges need to be addressed(23). Among these challenges, the integrity of the anatomy content is a primary concern due to various factors such as the quality of the original scan and the modeling design (24). Inaccuracies in the 3D models may result in misinterpretations of anatomical structures, which can be particularly problematic in medical education where precision is paramount. Another challenge is the amount of data required in creating a detailed and accurate representation of anatomical structures. The creation of high-resolution 3D models for XR anatomy education requires a fast-paced render system to handle heavy data while maintaining smooth performance for immersive interactivity and an optimal user experience. Consequently, the development of such content necessitates the adoption of specialized solutions and techniques to ensure seamless operation.

This research endeavors to propose optimal solutions and unique methodologies to the challenges encountered in XR anatomy education by developing a 3D Digital Twin (DT) of a Korean Standard Full-body Cadaver in metaverse. The efficacy of this innovation was assessed through a cohort of medical professionals and students from Ewha Woman’s University and 40 other nursing schools in Korea. The process of creating a 3D DT of a KSMC involved acquiring raw 3D object files for each bone using a 3D surface scanner. Subsequently, each 3D object file underwent refinement for visual accuracy while maintaining the integrity of the anatomical features of the original cadaver. This was achieved by adjusting the number of polygons to the optimal level using *Blender*, a 3D modeling software. The resulting model was then integrated into a metaverse platform built using Unity, which consisted of UX/UI elements designed to facilitate the immersive user experience. Furthermore, *Azure*, a cloud computing platform, was employed to provide real-time remote rendering services, which addressed the challenge of rendering high-resolution 3D models and alleviated the performance burden on the *HoloLens 2 AR HMD*. This component facilitated not only the individual user’s immersive experience but also allowed multiple users to interact and communicate with the digital twin cadaver in real-time, presenting a unique and innovative approach to anatomy education.

## MATERIALS AND METHODS

The process of creating a virtual 3D digital twin of a full-body Korean male cadaver involved collecting raw 3D object files for each bone using a 3D surface scanner. Subsequently, each 3D object file underwent refinement for visual accuracy while maintaining the integrity of the anatomical features of the original cadaver. This was achieved by adjusting the number of polygons to the optimal level using Blender, a 3D modeling software. The resulting model was then integrated into a metaverse platform built using Unity, which consisted of UX/UI elements designed to facilitate the immersive user experience. To enhance the interactivity and real-time communication capabilities of the metaverse platform, Microsoft Mixed Reality Toolkit (MRTK) and Photon Network were applied in Unity. Furthermore, Azure, Microsoft’s cloud computing platform, was employed to provide real-time remote rendering services, which addressed the challenge of rendering high-resolution 3D models and alleviated the performance burden on the HoloLens 2 AR HMD. This component not only facilitates the individual user’s immersive experience but also allows multiple users to interact and communicate with the digital twin cadaver in real-time, presenting a unique and innovative approach to anatomy education.

### From Cadaver to a 3D Digital Twin Cadaver

#### Collecting and Scanning

To create a 3D DT of a KSMC, we followed a multi-step process that began with the ethical procurement of a full-body cadaver from a certified Korean institution. A total of 206 individual bones of a male cadaver were donated to the Department of Anatomy of Catholic University of Korea and were then shared with Ewha Woman’s University for a full 3D scan.

For the 3D scanning process, we used a Leica BLK360 3D laser scanner, which offers several advantages over other 3D scanning methods. One of the main benefits of the BLK360 scanner is that it does not harm the cadaver, making it an ethical choice for capturing high-quality 3D scans. Additionally, the scanner is fast, accurate, and captures data with a high level of detail. To scan the cadaver body, we first placed it on a stable platform, and then used the BLK360 scanner to capture a series of 3D scans from multiple angles. These scans were then combined and processed using specialized software to create a complete and accurate 3D model of the cadaver.

The use of the Leica BLK360 scanner offers several advantages over other scanning methods. First, it provides high accuracy and resolution, which is essential for capturing the intricate details of the anatomy. Additionally, BLK360 scanner is a quick and efficient method, reducing the time required for the scanning process compared to other traditional methods. Most importantly, using this method allowed us to maintain the ethical standards of cadaver use, as the cadaver was kept intact and preserved for its intended purpose of donation.

#### Refining and Retopologizing the Mesh for Accuracy

After scanning the bones, each was then refined using 3D modeling softwares, called Blender and Maya. Refining a 3D scan in Blender or Maya is a crucial step as we develop anatomical content for the metaverse. This process involved using various tools and techniques to fill in any missing or incomplete parts of the model that may have been lost during the scanning process. To begin, the 3D scan is imported into Blender using the “Import” option in the “File” menu. Once imported, the model is carefully examined to identify any missing or incomplete parts using the “X-Ray” option in the “Viewport Overlays” menu.

Using the Sculpt Mode in Blender, the model can be manually sculpted and shaped to fill in any missing details or smooth out rough areas. Additionally, retopology techniques can be used to create a new, cleaner mesh over the existing mesh of the scan. This can make it easier to work with and modify, as well as provide a more accurate representation of the scanned object.

During the scanning process, it was discovered that some of the bones in the original cadaver exhibited synostosis, such as the middle and end phalanges of both fifth metatarsal bones, and the sacrum and coccyx. To address this issue, we consulted various medical resources and anatomical atlases as references and employed retopology techniques.

The fused bones were treated as single objects, and the six ear bones were combined with the 22 cranium bones for grouping purposes. After the mesh was retopologized, we utilized Mesh Modeling tools to further refine and shape the model. Specifically, these tools were used to add or subtract polygons, create new shapes, or modify existing ones, especially for extremely thin layers or deep holes in the bones that impeded proper 3D modeling.

#### Optimizing Polygons

In addition to refining the mesh of the 3D model, it is essential to optimize the number of polygons in the 3D models, especially for a real-time interactive experience. The number of polygons in a 3D model has a direct impact on its processing power and memory requirements for proper rendering. As the number of polygons increases, the performance of the virtual environment can significantly slow down. Therefore, it is crucial to optimize the number of polygons in 3D models to ensure smooth rendering and efficient performance in the metaverse.

We chose to optimize the number of polygons in the 3D objects to use techniques such as decimation, which involves reducing the number of polygons in a model while preserving its overall shape and structure. An important consideration when optimizing the number of polygons is to balance the level of detail required for specific use cases. Since this development creates anatomical content for educational purposes, it requires a higher level of detail and intricacy.

### Architecture for the metaverse platform to Facilitate a 3D digital **twin.**

#### HoloLens 2

HoloLens is a mixed-reality device that allows users to interact with holographic objects and digital content in real-world environments. Table 1 shows HoloLens 2 specifications. This provides a more immersive and engaging experience than traditional 2D screens or virtual reality devices. HoloLens transforms anatomical content into an immersive, dynamic experience, as users can manipulate the organs, bones, and any anatomical structures of the 3D DT with simple hand gestures and voice commands, allowing them to explore and learn at their own pace. HoloLens can also provide real-time feedback and guidance, enhancing the learning experience and providing a more hands-on approach to education.

**Table 1.**
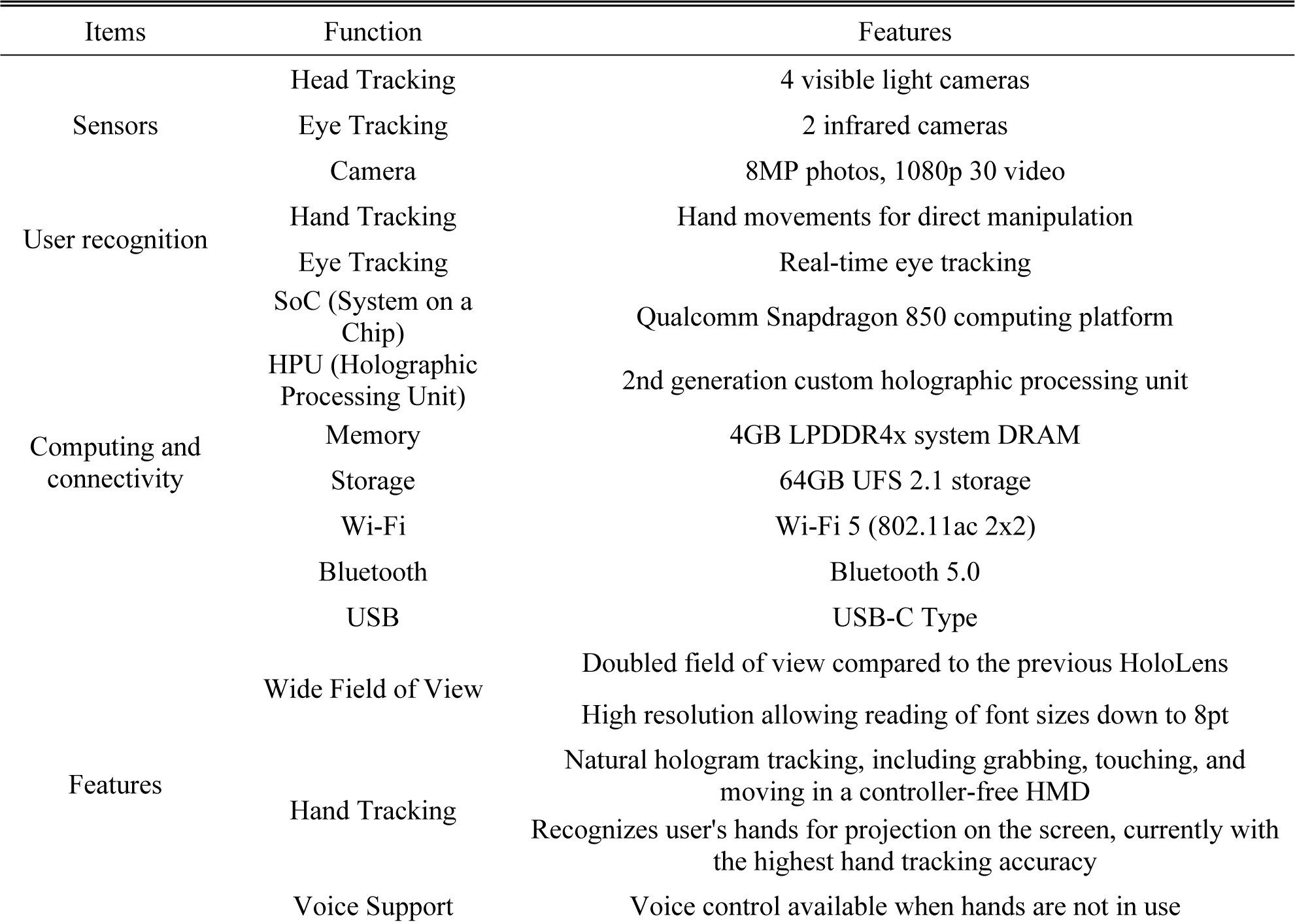
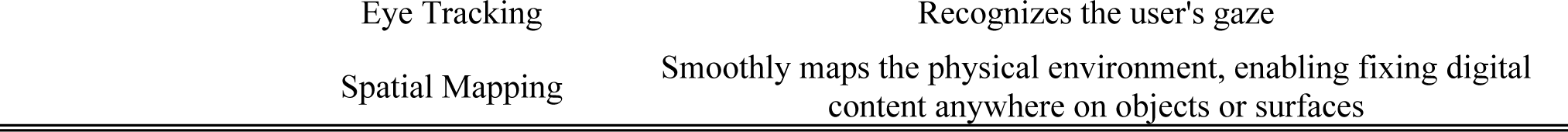
Specification of HoloLens 2.

#### Developing a Metaverse Platform and Adding Interactivity

Once the 3D models were complete, scripting is required to create an interactive model. As we utilized Unity, which is best compatible with the HoloLens2, and C# as it is a commonly used scripting language in Unity. By writing scripts, the model can be manipulated by the user. For example, the user can rotate the model or select a specific body part to receive additional information about it. In addition, we incorporated Mixed Reality Toolkit (MRTK), which is a collection of scripts, components, and assets that simplifies the development of mixed reality applications, to include pre-built components such as buttons and sliders, as well as tools for input management and spatial mapping. For additional functionalities such as real-time interaction among multiple users, we integrated Photon Network, a real-time multiplayer network engine designed for Unity. This integration adds to the collaborative learning experience and facilitates teaching scenarios. These tools provide essential components for creating realistic 3D DT models and simulations, enabling seamless mixed reality experiences and real-time collaboration and conference capabilities. As such, 3D DT models can be created with both realistic precision and computational efficiency.

### Development of XR contents

Remote rendering can be a creative way to develop Unity programs for high-quality 3D anatomy content, as it enables the rendering of complex and data-intensive 3D models without the need for expensive local hardware or extensive processing power.

With remote rendering, the 3D model is sent to a remote server, or a cloud-based service. The rendered images are then transmitted back to the user’s device, allowing smooth and high-quality visualization of the 3D anatomy content. This approach offers developers the advantage of ensuring optimal performance of their Unity programs across various devices, from low-powered laptops to high-end workstations. In addition, remote rendering can substantially cut the amount of the time and cost associated in developing and deploying Unity programs that feature data-intensive content like the interactive 3D DT models. By using cloud-based rendering services, developers can avoid the need for costly investments and focus on designing the most compelling interactive content for their users.

Overall, remote rendering is a powerful tool for developing Unity programs with smooth and high-quality 3D anatomy content. By leveraging the power of remote servers and cloud-based rendering services, developers can create immersive and engaging programs that provide a unique and valuable learning experience for students and professionals in the field of anatomy.

## RESULTS

### 3D digital twin cadaver

A 3D DT model, scanned from an actual human cadaver, underwent assembly modeling and was 3D printed. The production process utilized the internal filament removal method, a technique in 3D printing, to ensure both durability and weight. The model, with a size of 45 cm, was successfully printed, as shown in Fig 1 (a), and the assembled model is depicted in Fig 1 (b).

**Fig 1.**
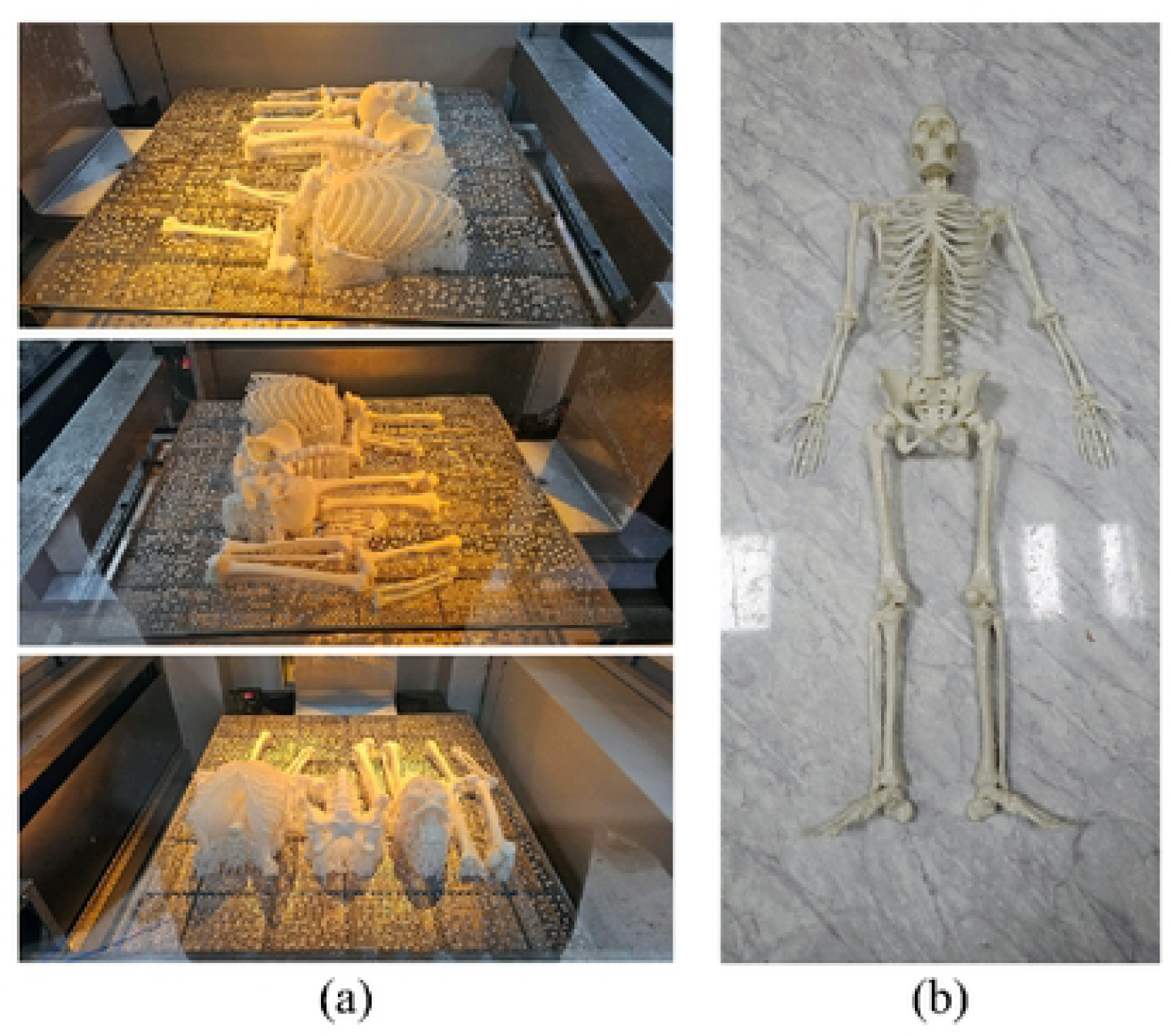
Bones printed using 3D printing: (a) The appearance of bones right after being 3D printed (b) Assembled result of the printed bones.

### Architecture for the metaverse platform

Within XR content, the utilization of the Photon Cloud network service platform is employed for inter-user data synchronization. In this content, the ‘REALTIME’ feature is utilized to real-time share the states of objects, and an ‘Addon Voice’ App ID key is obtained for the functionality of HoloLens 2 Voice. The acquired ID values are stored in the variables of the SharingServiceProfile script, and when the collaboration feature is initiated, the program automatically connects to the Photon Cloud server using the entered key values. Furthermore, in the Photon Provider configuration, the Provider, an object modularizing overall functionalities used after connecting to the Photon Cloud service, is concurrently configured with the connection to the remote rendering service. The initial launch screen of the metaverse platform offers features such as creating rooms for interaction with others or joining rooms established by other users.

The “Create Room” function, initiated by clicking the ‘Create Room’ button in the ‘Sharing Session Panel,’ follows the subsequent process: 1) It first verifies the presence of the Provider. Upon confirmation, it checks the count of objects created through the remote rendering service. 2) Upon identifying rendered objects, it removes all such objects and executes the Provider’s room creation function. 3) The room creation function of the Provider examines the names of all rooms created on the currently connected Photon server, and if no room with the same name exists, it utilizes the room creation function provided by the Photon SDK to generate a new room. The “Join Room” function employs the same process, using the same function as the room creation, and when selecting a room to join, it compares the set room name value with existing rooms. If a match is found, the user joins the specified room. Additionally, when the model position sharing feature is enabled, it interfaces with the Azure Anchor Service using the configured Id, Key, and Domain values. This ensures that remotely rendered models coexist in the same location, with the model’s reference point based on the person who created the room.

In metaverse, the functionality of remote rendering is available for mutual use. Upon creating a room and connecting to the Photon server, the current list of connected players is displayed on the right side. In the collaboration panel, the present connection status, room name, and the number of connected players are indicated.

### XR contents

#### Login and account creation

Table 2 presenting elements related to login and account creation. “User Login” delineates the functionality designed for user identification through login. Upon clicking the “Create Account” button, the HoloLens activates its internal Explore functionality, facilitating the transition to a linked Uniform Resource Locator (URL) address. Consequently, the registration window opens, allowing users to input the required data and complete the registration process.

**Table 2.**
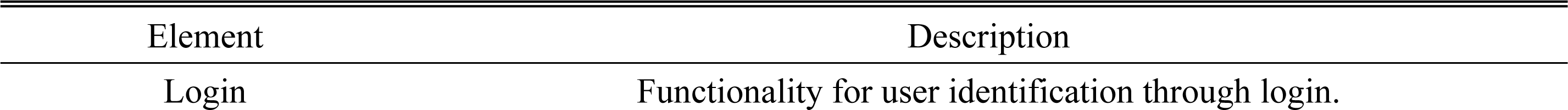
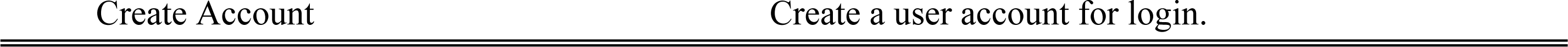
The functionality within the “User Login” section of XR content.

#### Menu

Table 3 presenting a compilation of menu and feature-related items. The “App Menu” incorporates diverse functionalities, including but not limited to tutorial, button selection, menu language change, tool function, partial palette function, description window function, collaboration function, and other menu options. The “Name” entry addresses the display of names based on language options. Fig 2 shows the menu screen of the manipulation tool. In the manipulation tool, users can move or detach bones to display them. Additionally, bones can be differentiated by changing colors, and there is a feature to output the surface of the bones if needed. Fig 3 shows an individual engaging with the XR content firsthand while donning the HoloLens.

**Table 3.**
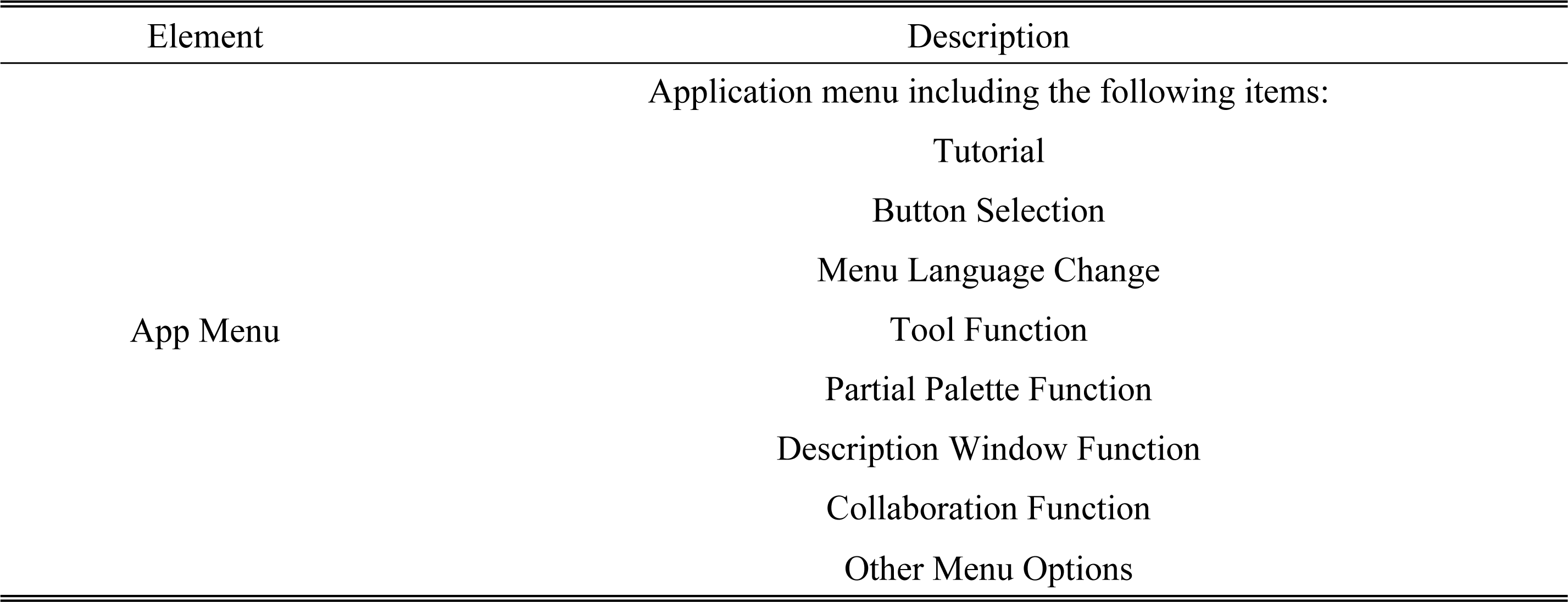
The functionality within the “App Menu” section of XR content.

**Fig 2.**
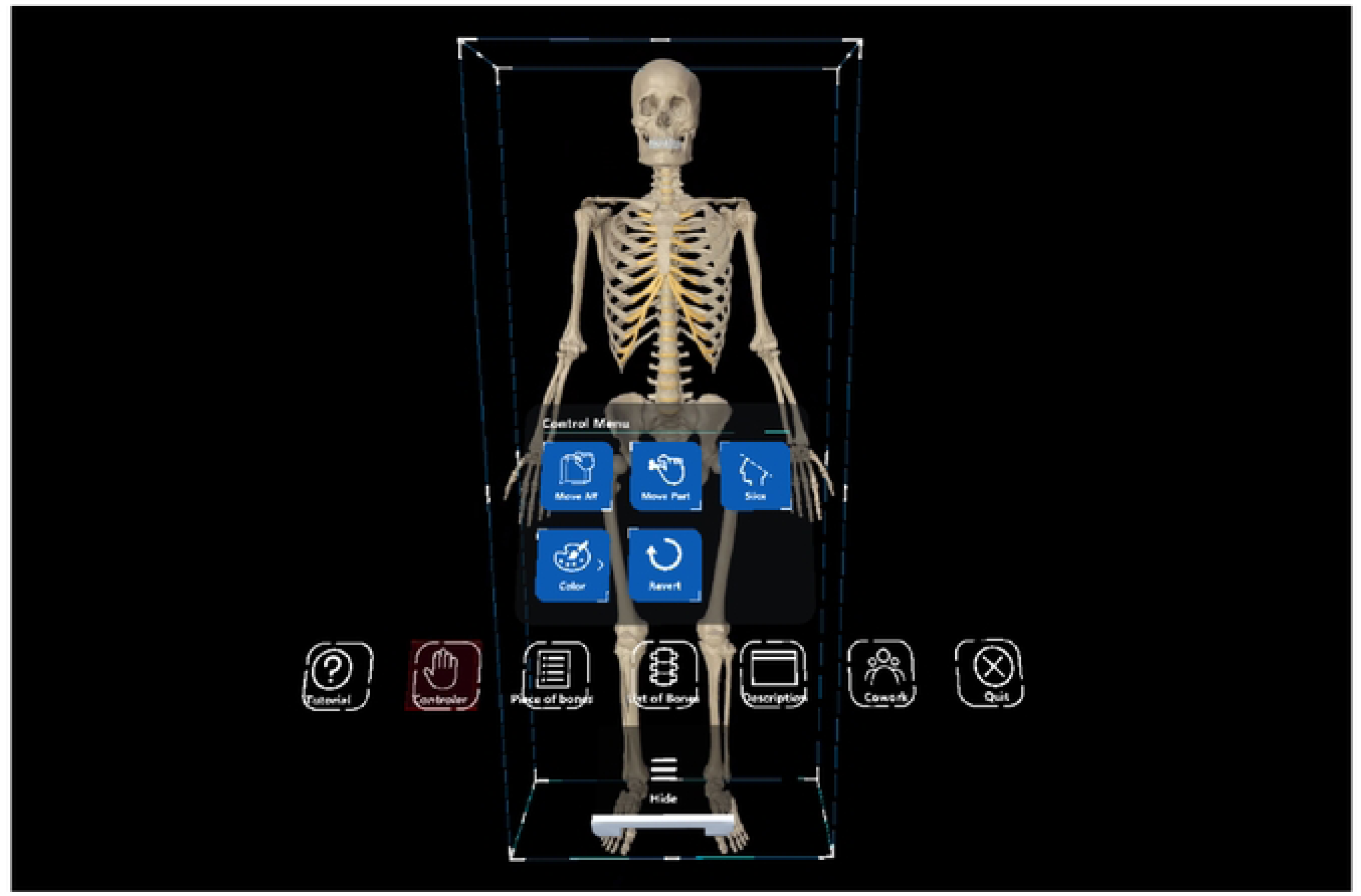
Menu screen of the manipulation tool. The manipulation menu includes functions to select, move, separate, and add colors to the desired bones.

**Fig 3.**
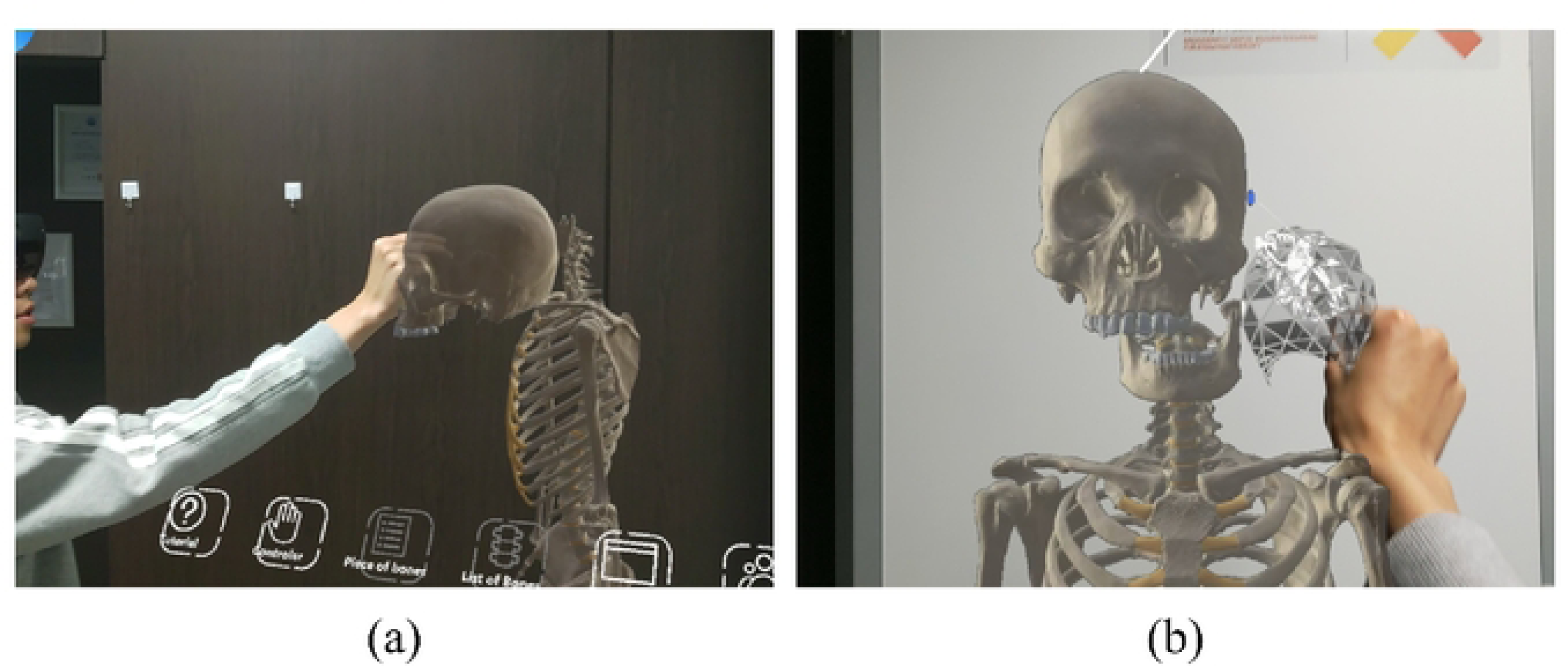
Menu screen of the manipulation tool. The manipulation menu includes functions to select, move, separate, and add colors to the desired bones. (a) Side view, (b) Front view.

#### Functions

Table 4 presenting elements related to tool functions, partial palette, description window, and collaboration function. “Tool Functions” encompasses a range of functionalities such as moving the entire model, moving individual parts, cross-section cutting, changing color, and undo function. “Partial Palette” covers various functionalities related to specific body areas. The “Description Window” provides detailed images and descriptions for items within the partial palette. Finally, “Collaboration Function” includes features like creating a room, joining a room, sharing or canceling model location, and ending or exiting the collaboration session.

**Table 4.**
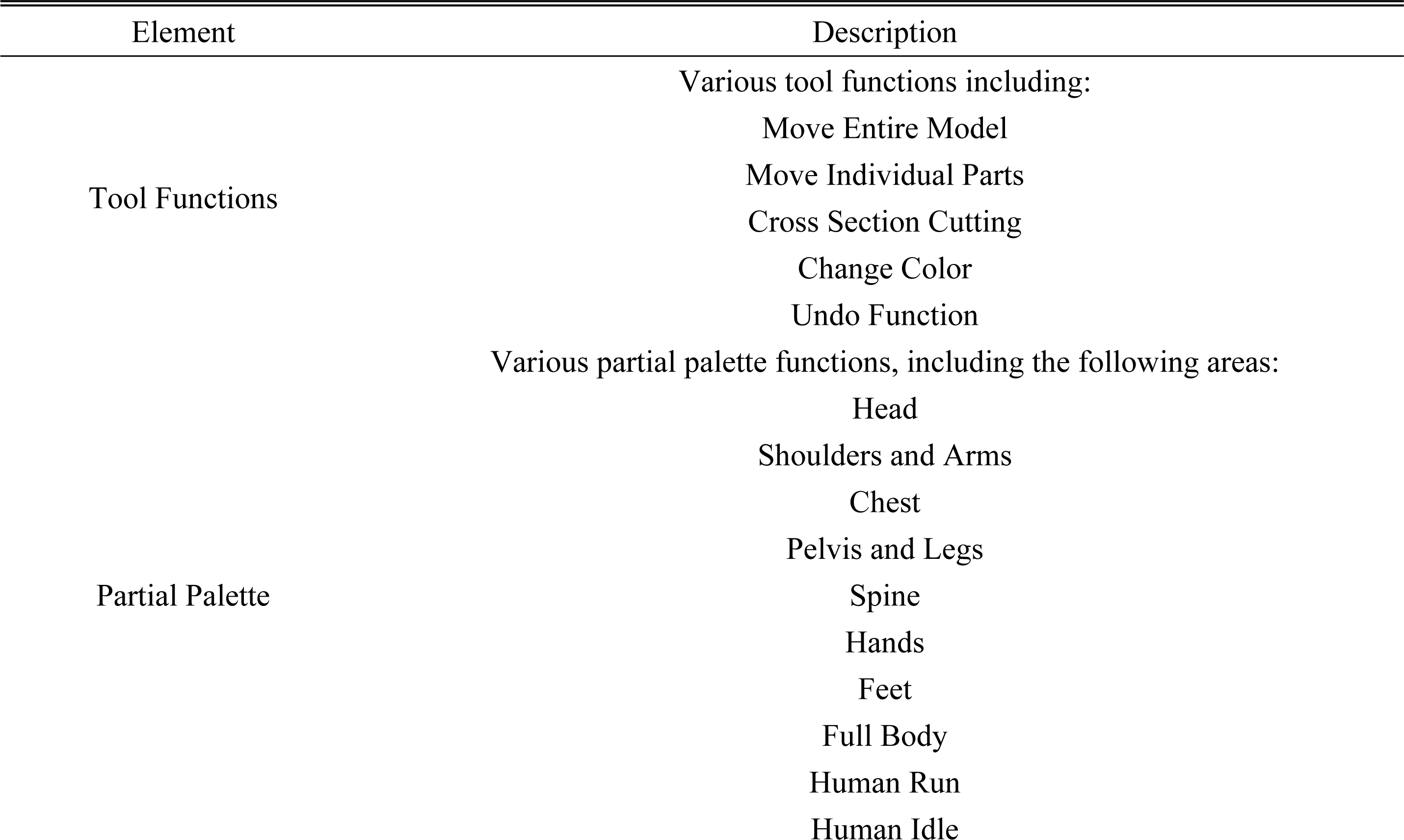
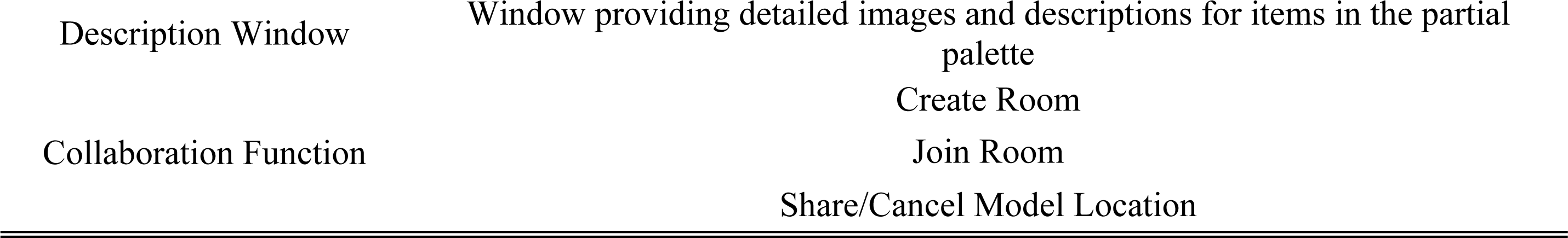
Various elements incorporated in XR content.

## DISCUSSION

This research introduces a novel advancement in conventional anatomy education, addressing numerous challenges posed by traditional cadaver dissection. The utilization of 3D scanning and modeling techniques results in the creation of high-fidelity interactive 3D DT models with precise anatomical information for deeper learning experiences. The incorporation of technologies such as Photon Network facilitates real-time collaboration among students, fostering deeper engagement, and facilitating a collaborative educational environment. Additionally, the adoption of cloud-based remote rendering technology ensures seamless and high-quality visualization of 3D models across diverse devices, contributes to cost reduction in hardware and enhancing accessibility.

The final product of this novel development provides valuable insight towards the future of anatomy education, highlighting the pivotal role of XR technology and 3D DTs in offering boundless learning opportunities for users. Ethical considerations related to the development and use of 3D digital twins are emphasized, addressing concerns associated with practical anatomy education and underscoring a commitment to upholding ethical standards.

The results presents the need for improvement and scalability, with the technology having the potential to expand and encompass more anatomical content, finding application in various medical fields and offering a versatile educational platform. XR technology, including the metaverse, emerges as a transformative force in anatomy education, suggesting potential replacements for traditional cadaver dissection, enhanced accessibility, and efficient learning experiences, particularly through devices like HoloLens 2.

Amidst the high performance of this novel development, several areas for improvement were found. As shown in Fig 4 (a), the length and thickness of the sutures at the junctions between the lumbar vertebrae and the torso, as well as between the torso and the head, were found to be insufficient and required reinforcement. Although there were no issues with support, there was a tendency for swaying due to the weight distribution between the torso and the head bones. Next, as depicted in Fig 4 (b), there was a need to change the material of the rib cartilage to reduce the risk of damage. To achieve a connection between different materials, the model will be reworked using a suturing method. Lastly, as illustrated in Fig 4 (c), there was a need to address the unstable structure where the wing bones were only connected to the chest bones. To improve this aspect in the future, additional connection points between the rib bones and the wing bones will be incorporated.

**Fig 4.**
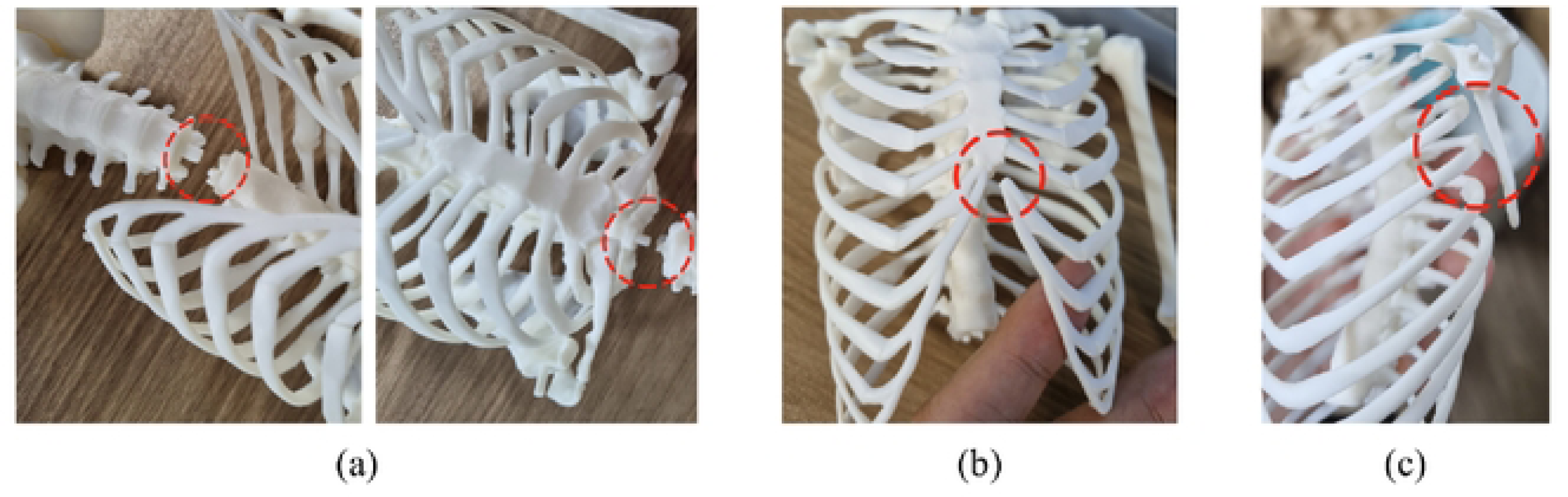
The content screen corresponding to the “Partial Palette” in the “App Menu.” “Partial Palette” allows the selection of specific bone regions and provides options to observe the movements of the human body.

The model’s junctions include the skull-neck, neck-torso, shoulder-upper arm, upper arm-lower arm, lower arm-wrist, torso-lumbar vertebrae, lumbar vertebrae-thigh bone, thigh bone-shin bone, shin bone-ankle bone, etc., as demonstrated in Fig 5. The detailed enhancement plan for the 3D skeletal cadaver is outlined as follows.

**Fig 5.**
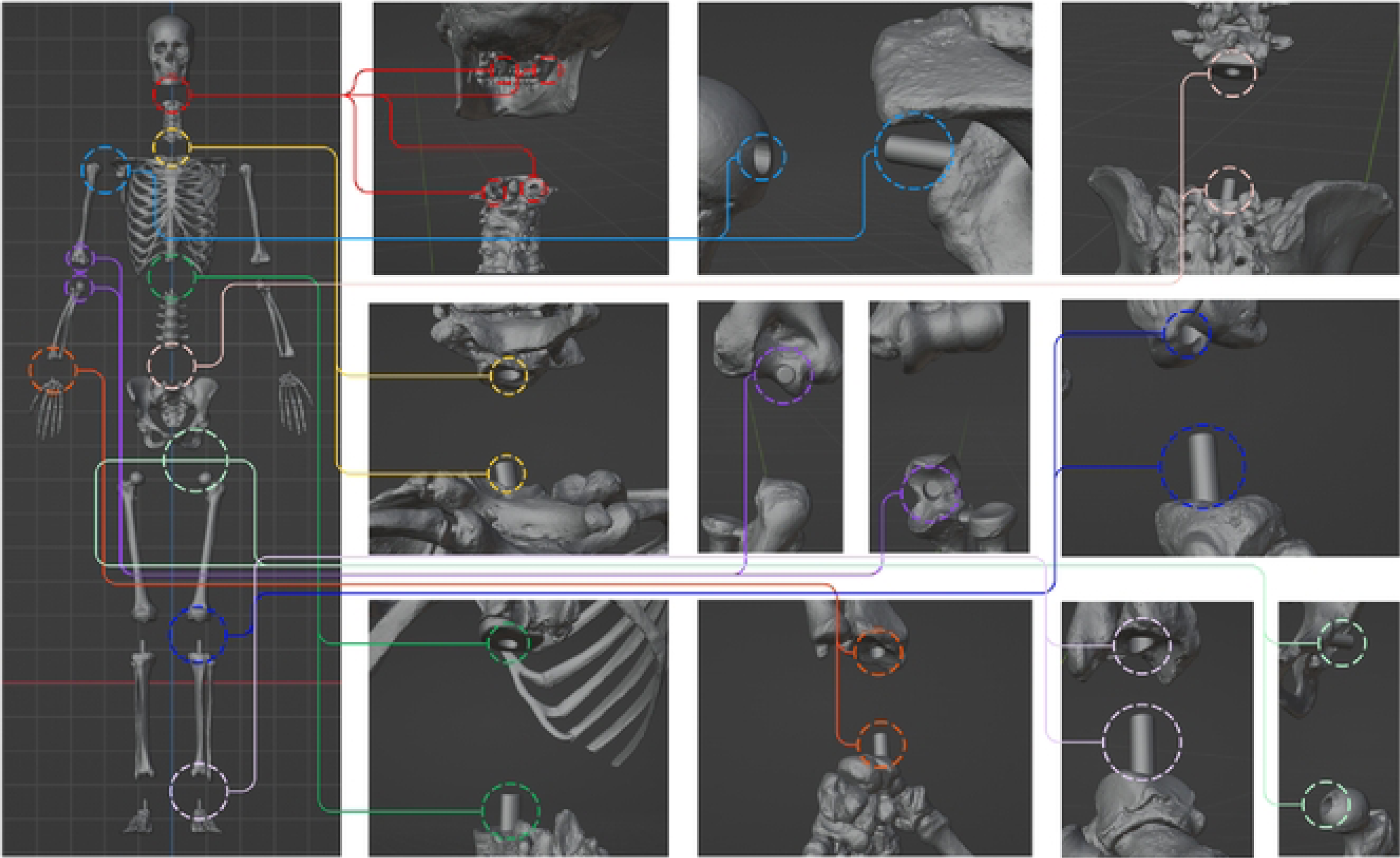
Modeling of articulations for the creation of a 3D skeletal model.

The exploration of XR anatomy education revealed specific technical challenges, which were manifested by the process bone scanning and modeling processes. In addition, while such novel XR educational tools signify a significant advancement in medical education, challenges such as battery life, high costs, and internet connectivity persist. Nonetheless, these developments mark a substantial leap towards a new era in collaborative and interactive learning, ensuring improved performance and user experience. They not only facilitate the creation of interactive 3D DTs from MRI or CT images, expanding beyond gross anatomy to specialized curriculums, but also open opportunities in integration with artificial intelligence to enhance learning content based on data-driven insights and user experiences. The positive outcomes underscore the promising role of XR technology in healthcare and advocate for further innovative applications.

## CONCLUSION

Technological innovation in the field of anatomy education is poised to play a crucial role in shaping the future generation of medical and healthcare professionals. The proposed content in this study holds significance in overcoming the limitations of traditional anatomy education and spearheading innovation in the learning experiences of future practitioners. Sustained efforts in the development and enhancement of such technologies are imperative, as they are poised to furnish learners with essential tools for success in the evolving landscape of healthcare in the future.

## ACKNOWLEDGEMENTS

This research supported by the Korea Institute of Energy Technology Evaluation and Planning (KETEP) grant was funded by the Korea government (MOTIE) (20227410100050) and supported by the Basic Science Research Program through the National Research Foundation of Korea (NRF) funded by the Ministry of Education (grant number NRF-2022R1H1A2092091) and this work was supported by the Technology development Program(RS-2023-00257618) funded by the Ministry of SMEs and Startups(MSS, Korea).

## AUTHOR CONTRIBUTIONS

**Conceptualization:** So Hyun Ahn, Seung Ho Han

**Formal analysis:** Dong Hyeok Choi, Seo Yi Choi

**Resources:** Rena Lee, So Hyun Ahn, Sung Ho Cho, Seung Ho Han

**Writing – original draft:** Dong Hyeok Choi, Seo Yi Choi

**Writing – review & editing:** Dong Hyeok Choi, Seo Yi Choi, Rena Lee, So Hyun Ahn, Sung Ho Cho, Seung Ho Han

## REFERENCE

1. Böckers A, Jerg-Bretzke L, Lamp C, Brinkmann A, Traue HC, Böckers TM. The gross anatomy course: An analysis of its importance. Anatomical sciences education. 2010;3(1):3–11.

2. Habicht JL, Kiessling C, Winkelmann A. Bodies for anatomy education in medical schools: an overview of the sources of cadavers worldwide. Academic Medicine. 2018;93(9):1293.

3. Tapia-Nañez M, Quiroga-Garza A, Guerrero-Mendivil FD, Salinas-Alvarez Y, Jacobo-Baca G, De la Fuente-Villarreal D, et al. A review of the importance of research in Anatomy, an evidence-based science. Eur J Anat. 2022;26(4):477–86.

4. Štrkalj G, Pather N. Beyond the sex binary: Toward the inclusive anatomical sciences education. Anatomical sciences education. 2021;14(4):513–8.

5. Shahrvini B, Baxter SL, Coffey CS, MacDonald BV, Lander L. Pre-clinical remote undergraduate medical education during the COVID-19 pandemic: a survey study. BMC Medical education. 2021;21(1):1–13.

6. McLachlan JC, Bligh J, Bradley P, Searle J. Teaching anatomy without cadavers. Medical education. 2004;38(4):418–24.

7. Varner C, Dixon L, Simons MC. The past, present, and future: a discussion of cadaver use in medical and veterinary education. Frontiers in Veterinary Science. 2021;8:720740.

8. Shapiro L, Bell K, Dhas K, Branson T, Louw G, Keenan ID. Focused multisensory anatomy observation and drawing for enhancing social learning and three-dimensional spatial understanding. Anatomical Sciences Education. 2020;13(4):488–503.

9. Attardi SM, Harmon DJ, Barremkala M, Bentley DC, Brown KM, Dennis JF, et al. An analysis of anatomy education before and during Covid-19: August–December 2020. Anatomical sciences education. 2022;15(1):5–26.

10. Amiralli H, Joseph S. Dissecting the future: A critical review of anatomy’s past, present, and future following the carnegie foundation’s call for medical education reform. Journal of The Anatomical Society of India. 2019;68(4):306-.

11. Papapanou M, Routsi E, Tsamakis K, Fotis L, Marinos G, Lidoriki I, et al. Medical education challenges and innovations during COVID-19 pandemic. Postgraduate medical journal. 2022;98(1159):321-7.

12. Zhang J, Lu Q, Shi L. The influence of telemedicine on capacity development in public primary hospitals in China: a scoping review. Clinical eHealth. 2022.

13. Calton B, Abedini N, Fratkin M. Telemedicine in the time of coronavirus. Journal of pain and symptom management. 2020;60(1):e12–e4.

14. Alkhowailed MS, Rasheed Z, Shariq A, Elzainy A, El Sadik A, Alkhamiss A, et al. Digitalization plan in medical education during COVID-19 lockdown. Informatics in medicine unlocked. 2020;20:100432.

15. Moro C, Birt J, Stromberga Z, Phelps C, Clark J, Glasziou P, et al. Virtual and augmented reality enhancements to medical and science student physiology and anatomy test performance: A systematic review and meta-analysis. Anatomical sciences education. 2021;14(3):368–76.

16. Taylor L, Dyer T, Al-Azzawi M, Smith C, Nzeako O, Shah Z. Extended reality anatomy undergraduate teaching: A literature review on an alternative method of learning. Annals of Anatomy-Anatomischer Anzeiger. 2022;239:151817.

17. Harmon DJ, Attardi SM, Waite JG, Topp KS, Smoot BJ, Farkas GJ. Predictive factors of academic success in neuromusculoskeletal anatomy among doctor of physical therapy students. Anatomical Sciences Education. 2023;16(2):323–33.

18. Richards S. Student Engagement Using HoloLens Mixed-Reality Technology in Human Anatomy Laboratories for Osteopathic Medical Students: An Instructional Model. Medical Science Educator. 2023;33(1):223–31.

19. Koo H. Training in lung cancer surgery through the metaverse, including extended reality, in the smart operating room of Seoul National University Bundang Hospital, Korea. Journal of educational evaluation for health professions. 2021;18.

20. Pottle J. Virtual reality and the transformation of medical education. Future healthcare journal. 2019;6(3):181.

21. Jones D, Hazelton M, Evans DJ, Pento V, See ZS, Van Leugenhaege L, et al. The road to birth: Using digital technology to visualise pregnancy anatomy. Digital Anatomy: Applications of Virtual, Mixed and Augmented Reality: Springer; 2021. p. 325–42.

22. Sandrone S. Medical education in the metaverse. Nature Medicine. 2022;28(12):2456–7.

23. Papa V, Vaccarezza M. Teaching anatomy in the XXI century: new aspects and pitfalls. The Scientific World Journal. 2013;2013.

24. Chickness JP, Trautman-Buckley KM, Evey K, Labranche L. Novel development of a 3D digital mediastinum model for anatomy education. Translational Research in Anatomy. 2022;26:100158.

